# Neuronal-Activity-Related Sodium (NARS) fMRI Reveals Millisecond Neuronal Dynamics Beyond Hemodynamic Readouts

**DOI:** 10.64898/2026.02.09.704765

**Authors:** Xin Yu, Xiaochen Liu, Grace Yu, Yuanyuan Jiang, Nivetha Pasupathy, David Hike, Xiaoqing Alice Zhou

**Affiliations:** Translational Neuroimaging and Neural Control Laboratory, Martinos Center for Biomedical Imaging, MGH/HMS

**Keywords:** ultra-fast fMRI, sodium-23 MRI, quadrupolar relaxation, functional neuroimaging, non-hemodynamic contrast, neuronal activity

## Abstract

Brain-wide, noninvasive methods that directly resolve neuronal activity on millisecond timescales are still lacking in neuroimaging. Hemodynamic fMRI (^1^H-based) provides whole-brain maps of activity, yet its vascular origin creates spatial and temporal displacements from neuronal events that complicate interpretation, especially in disease conditions where neurovascular coupling is altered. Here, we developed an ultrafast ^23^Na fMRI platform at 14 T that combines a reshuffled k-t 3D gradient echo readout (TR/TE, 10ms/1ms) with an implantable RF coil and respiration-gated acquisition. This configuration provides the sampling rate and SNR needed to probe quadrupolar T2*-weighted single quantum sodium dynamics on the neuronal timescale. Across rats and mice, somatosensory forepaw stimulation produced a localized ^23^Na signal decrease of ~2-3% in the FP-S1, peaking ~10-30 ms post-stimulus. The activity pattern is well matched with conventional BOLD-fMRI maps acquired in the same animals. Trial-by-trial measurements during simultaneous iGluSnFR glutamate fiber photometry demonstrated that larger evoked glutamate transients coincided with larger NARS decreases, supporting a neuronal origin of the NARS contrast. We interpret the negative NARS response as a transient activity-dependent redistribution of sodium ions toward restricted, protein-rich microdomains, where more restricted rotational dynamics may accelerate T2*short decay and produce a 2-3% signal decrease without requiring large changes in bulk sodium concentration. Together, these results establish neural activity-related sodium (NARS) fMRI as a viable approach for direct, mesoscale neuronal mapping with MRI at millisecond resolution.

**Graphical Abstract:** A reshuffled k-t 3D gradient echo (GRE) readout with an implantable figure-8 RF coil and respiration-gated trials captures a ~2-3% negative ^23^Na response peaking 10-30 ms after forepaw stimulation in the FP-S1. NARS fMRI responses co-localize with BOLD-fMRI and scale with iGluSnFR glutamate transients, consistent with activity-dependent shifts toward short-T_2_* sodium microenvironments.

**Table of Contents (TOC) Blurb:** Yu et al. introduce NARS-fMRI, an ultrafast ^23^Na method that resolves millisecond neuronal dynamics in rodent FP-S1. The negative sodium response (10-30 ms) co-localizes with BOLD and correlates with glutamate photometry, supporting a direct neuronal origin.

## Introduction

Hemodynamic functional MRI (fMRI) maps brain activity by detecting changes in the magnetic relaxation of water proton (^1^H) spins driven by alterations in cerebral blood volume, blood flow, and oxygenation within activated regions^1–5^. Because the measured spins originate from vascular compartments rather than neurons, hemodynamic fMRI inherently introduces spatial and temporal dislocation from the true neuronal sources^6–11^. Moreover, the signals reflect an indirect cascade of neurovascular coupling, integrating not only neuronal activity but also autonomic regulation and intrinsic vascular dynamics^12–15^. These features create substantial challenges for clinical and translational applications, particularly in diseased brains where blood flow, vascular reactivity, and metabolism can deviate from neuronal activity. Thus, directly measuring neuronal activity via NMR signals remains a central unmet goal in neuroimaging. To approach a “true fMRI” contrast that is directly linked to neuronal processes, such as action potentials (APs) or local field potentials (LFPs) driven by transmembrane ion flux, x-nuclei MRI offers a promising avenue with the potential to overcome the fundamental limitations of proton(^1^H)-based hemodynamic fMRI.

Sodium is the key ion driving neuronal excitability, playing an essential role in the generation and propagation of APs^16–18^. The sodium NMR signal, particularly the quadrupolar relaxation behavior of the ^23^Na nucleus (spin-3/2), has been extensively characterized across diverse microenvironments, from solid-state systems to the living brain^19–24^. Building on this foundation, numerous studies have attempted to obtain semi-quantitative *in vivo* measurements of intracellular and extracellular ^23^Na compartments^25–27^. However, the inherently low signal-to-noise ratio (SNR) of ^23^Na NMR continues to limit its achievable spatial and temporal resolution compared with proton-based anatomical and functional MRI. Despite these constraints, several groups have demonstrated the feasibility of *in vivo* ^23^Na mapping in rodent brains by employing massive signal averaging on ultra-high-field scanners (9.4 T to 21 T) ^28–32^. Additional work has examined ^23^Na-dependent cerebral blood flow dynamics in rats^33^, and potential neurovascular-coupled ^23^Na signal changes have been reported in humans using modified ultra-short-echo (UTE) sequences^34^. Yet, a major technical barrier remains in how to capture dynamic ^23^Na signal fluctuations with the millisecond-level temporal precision required to resolve neuronal events. Overcoming this limitation is essential for establishing neuronal activity-related sodium (NARS)-fMRI as a direct and physiologically grounded alternative to conventional hemodynamic fMRI.

A central challenge in developing functional ^23^Na-based MRI is achieving sufficient SNR for detecting the inherently weak ^23^Na-NMR signal, which is estimated to be 1/20,000–1/30,000 of the *in vivo* proton (^1^H) signal due to its lower NMR sensitivity (^1^H:^23^Na ≈ 1:0.09) and markedly lower physiological concentration (~40 mM for ^23^Na vs. ~110 M for ^1^H)^25^. Bridging this gap requires advances that jointly improve temporal encoding efficiency and local detection sensitivity while preserving ultra-fast imaging capability. Two complementary technical advances provide a pathway toward this goal. Two complementary technical developments provide this foundation. First, reshuffled k–t space gradient-echo acquisitions introduced a new strategy for temporal encoding by interleaving k-space sampling with the stimulation paradigm, enabling laminar- and vascular-specific functional mapping^35–38^. The resulting ultra-short repetition times (TR=5-10 ms) permit high-temporal-resolution sampling of T_2_*-weighted signals^39,40^. Beyond accelerating acquisition, this framework has enabled single-vessel mapping of BOLD and CBV responses ^35,41^, revealing rapid microvascular dynamics consistent with established models of microvascular contributions to BOLD signal specificity^15,42–44^ and with high-resolution laminar fMRI studies at ultra-high field^45,46^. Second, advances in implantable RF coil technology have transformed local detection sensitivity by positioning miniature detectors directly adjacent to the tissue of interest ^47–50^. These optimized coils have enabled real-time, ultra-high-resolution functional imaging across multiple contrast mechanisms, ranging from balanced steady-state free precession(bSSFP)^51,52^, UTE sequences^53^, and even echo-planar imaging (EPI) at nearly 100 μm isotropic resolution in rodents^54,55^, reaching a mesoscale imaging regime^13^ that conceptually parallels submillimeter-resolution human brain functional mapping with next-generation ultra-high-field systems^56–60^. Together, reshuffled temporal encoding and localized RF detection represent converging advances that substantially narrow the intrinsic sensitivity gap of ^23^Na MRI, establishing the experimental framework necessary to approach neuronal activity–related sodium (NARS) fMRI at 14 T.

In the present study, we applied a reshuffled k–t space gradient-echo sequence to map evoked NARS-fMRI responses in the forepaw primary somatosensory cortex (FP-S1) of anesthetized mice and rats at 14 T. Here, we detected a robust ^23^Na signal decrease of ~2-3% in FP-S1 occurring 10-30 ms after electrical forepaw stimulation. To further validate the neuronal origin of this signal, we performed simultaneous NARS-fMRI and fiber-photometry recordings of glutamate (Glu) dynamics in the FP-S1 using the genetically encoded indicator iGluSnFR as a surrogate for LFPs ^61^. The strong correlation between evoked glutamate transients and the ^23^Na signal drop provides compelling evidence for a direct neuronal contribution to the observed NARS-fMRI contrast. Together, these findings establish NARS-fMRI as a viable method for direct neuronal activity mapping with MRI.

## Results

### Establishing ultra-fast NARS-fMRI at 14 T

To directly probe sodium signal dynamics at the neuronal timescale, we integrated an implantable figure-8 RF coil with a reshuffled k-t 3D gradient-echo readout (TR/TE at 10 ms/1 ms), enabling 100 Hz sampling of 23Na T2*-weighted images (Fig. 1A, B). The coil is rigidly affixed above the skull, providing stable B1 loading, bilateral somatosensory coverage, and a substantial SNR gain given the proximity between brain and external loop detector (Fig. 1C). The reshuffled acquisition interleaves k-space lines with the stimulation paradigm, allowing reconstruction of rapidly evolving sodium signals within 300 ms stimulus windows while with altered phase-encoding steps across repetitions. These design explicitly leverage the short T_1_ of ^23^Na at 14 T (~35–50 ms) and its biexponential quadrupolar T2* (T2*_short_ ≈ 0.5-5 ms) ^20,22,62–65^, which together favor single-quantum ^23^Na T2* weighted measurements with a gradient-echo approach. The *in vivo* T1 and T2* characterization at the cortex (Supp. Fig. 1) empirically supports these sequence choices, confirming compatibility with TE = 1 ms and rapid repetition.

**Figure 1:**
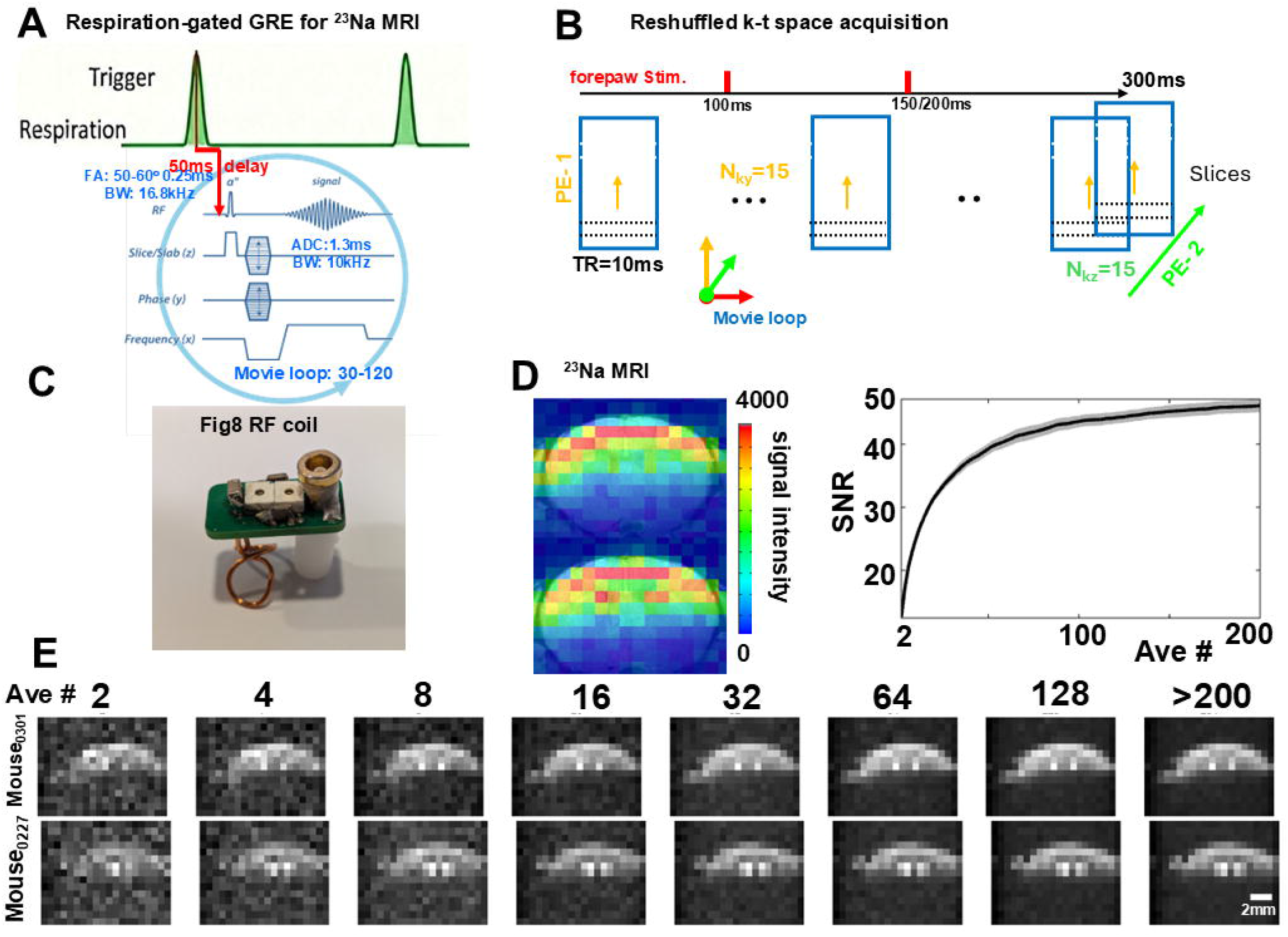
Establishing ultrafast NARS-fMRI at 14 T. (A). The timing diagram of the 23Na-based GRE sequence illustrating respiration-gated movie-loop acquisition within a single respiratory cycle (spoiler gradients were applied to suppress residual transverse coherences.). (B). The schematic of the reshuffled k–t space 3D gradient-echo (GRE) acquisition used for NARS-fMRI. Phase-encoding steps are interleaved across stimulus-locked movie loops, enabling ultrafast sampling (TR = 10 ms, TE = 1 ms) of dynamic ^23^Na T_2_*-weighted signals. (C) Photograph of the implantable figure-8 ^23^Na RF coil. (D) Quantification of cortical ^23^Na SNR as a function of averaging, demonstrating convergence to SNR ≈ 45 after ~200 repetitions (mean±s.e.m., n=5). Left panel is the color-coded ^23^Na magnitude images overlapped on FLASH anatomical ^1^H MRI images. (E) Representative ^23^Na magnitude images reconstructed with increasing numbers of averaged movie loops, illustrating progressive SNR improvement.

To contextualize feasibility based on the prior ultra-high resolution ^1^H-based fMRI studies with 14T^51,52,54,55^, we estimated an effective narrowing of the intrinsic ^23^Na/^1^H SNR gap by combining several experimentally grounded factors. First, NARS-fMRI uses larger voxel dimensions (600 × 600 × 1200 µm^3^ for rats, or 600 × 600 × 800 µm^3^ for mice) compared with ~100 µm isotropic resolution typical of CBV-fMRI, yielding a volumetric gain of ~432:1 or ~288:1^66^. Second, the figure-8 implantable coil (4-5 mm for each loop) provides an approximate two-fold sensitivity advantage relative to single-loop (~10 mm) surface coils^55^, Third, we considered the temporal sampling advantage enabled by the distinct relaxation properties and physiological timescales of sodium imaging. Conventional hemodynamic fMRI typically employs ~30 s stimulation epochs with TR ≈ 1-2 s due to the sluggish vascular response and long brain T_1_ (~3-4 s). In contrast, the NARS-fMRI paradigm targets millisecond-scale neuronal dynamics; with a ^23^Na T_1_ of ~30-40 ms at 14 T, ultrashort TR sampling (~10 ms) can be used within a ~300 ms acquisition window. Consequently, for a time span equivalent to a single 30 s hemodynamic epoch, approximately one hundred ultrafast sodium epochs can be acquired, yielding an estimated ~10-fold temporal SNR gain through repetitive averaging. Together, these factors suggest an overall ~6000-8000 fold effective SNR enhancement relative to the intrinsic ^23^Na-to-^1^H sensitivity difference, reducing the nominal gap from ~1/30,000 to approximately ~1:5 at the reconstructed image level. Consistent with this estimate, Fig. 1D and E show that the ^23^Na MRI SNR increases with the number of averages and approaches ~45 after less than 200 repetitions.

To minimize non-physiological signal dynamics, we combined respiration-gated windows with ultrafast ^23^Na readouts. Gating each stimulation epoch (e.g., 300 ms for mice) to a single respiratory cycle suppressed the aliasing artifact of the respiration-induced B_₀_ offsets in the looped ultrafast time series. In parallel, the variable inter-epoch intervals inherent to the respiratory rhythm (Suppl Fig. 2) promoted near-full T_1_ recovery and reduced short-TR spoiling-related steady-state oscillations.^67^, limiting sequence-induced intensity transients to a brief, reproducible first-frames saturation at the start of each window (Supp. Fig. 3). This dual control yields a stable baseline for detecting tens-of-milliseconds NARS-fMRI responses.

### Mapping FP-S1 activation in rats and mice with NARS-fMRI and BOLD-fMRI

We mapped the FP-S1 activation using NARS-fMRI and BOLD-fMRI in the same animals. In rats (Fig. 2) and mice (Fig. 3), statistical β-coefficient maps show co-localized activation in the FP-S1 across modalities. NARS-fMRI highlights a focal negative ^23^Na response, whereas BOLD-fMRI shows the canonical positive hemodynamic response in the same slices. There is strong spatial correspondence between the NARS-identified cortex and the BOLD-defined bilateral FP-S1. Time courses extracted from activated FP-S1 demonstrate a rapid ^23^Na signal decrease peaking ~10-30 ms after forepaw stimulation, which is consistent with the evoked LFP timescale. Across experiments, the evoked ^23^Na response is ~2-3% and localized to FP-S1of both rats and mice (Figs. 2, 3).

**Figure 2:**
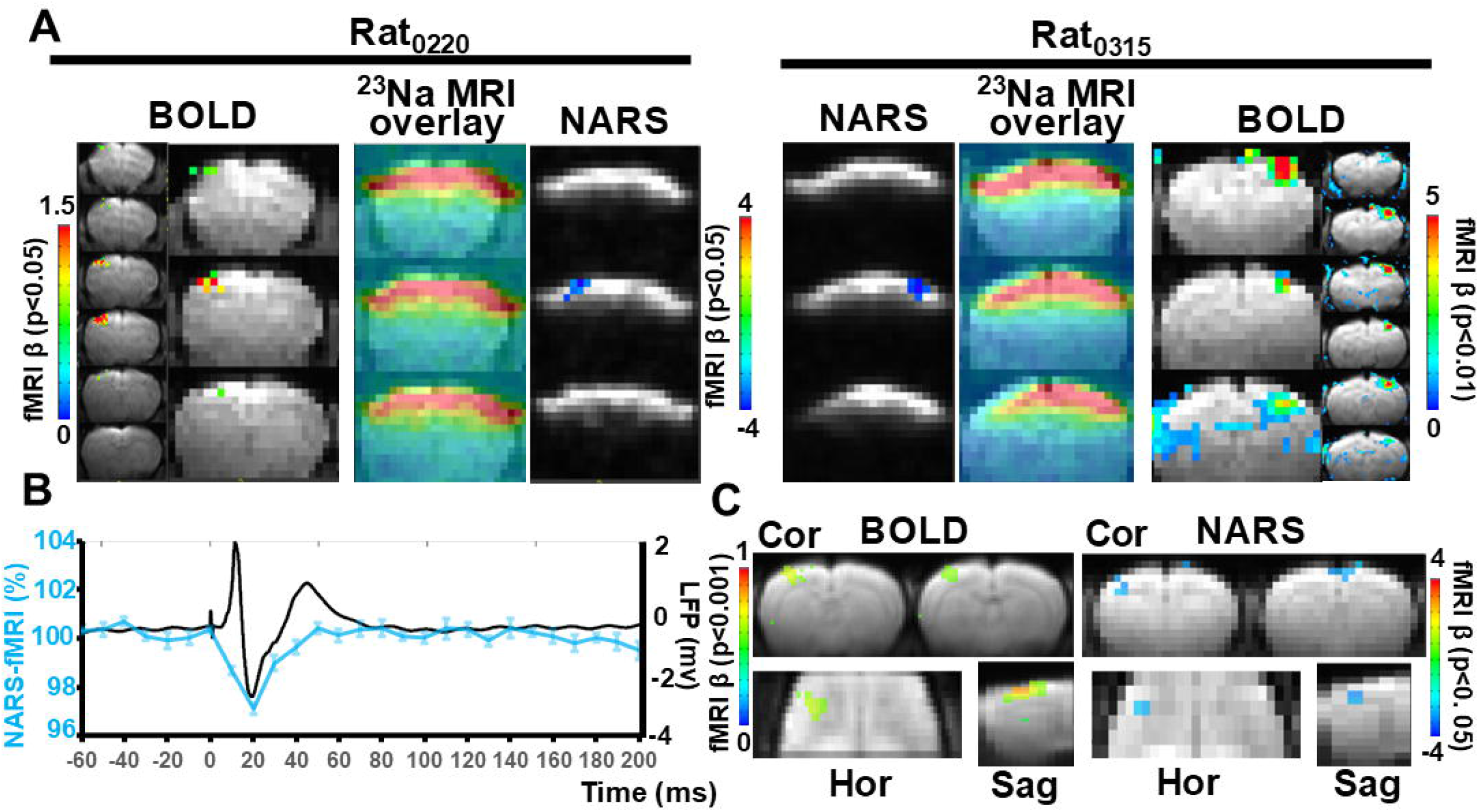
NARS-fMRI maps FP-S1 activation in rats. (A). BOLD and NARS-fMRI functional maps at different hemispheres from two representative rats. Left panel presents activation at the FP-S1 of the left hemisphere. Right panel presents activation at the FP-S1 of the right hemisphere. Positive BOLD activation is shown in statistical β-coefficient maps with in-plane 100×100µm^2^ and smoothed resolution matching ^23^Na images. The color-coded 23Na MRI images overlapped on the ^1^H EPI images. NARS-fMRI showing a focal negative ^23^Na response in the corresponding activated FP-S1 detected with BOLD. (B). Time courses extracted from FP-S1 showing a rapid NARS decrease peaking ~10-30 ms after stimulus onset (mean±s.e.m, n=10), compared with the evoked LFP spikes at a similar time-scale. (C) Group-averaged NARS-fMRI response amplitude across rats, showing a consistent ~2-3% negative signal change, which is well spatially aligned with BOLD activation maps (n=10).

**Figure 3:**
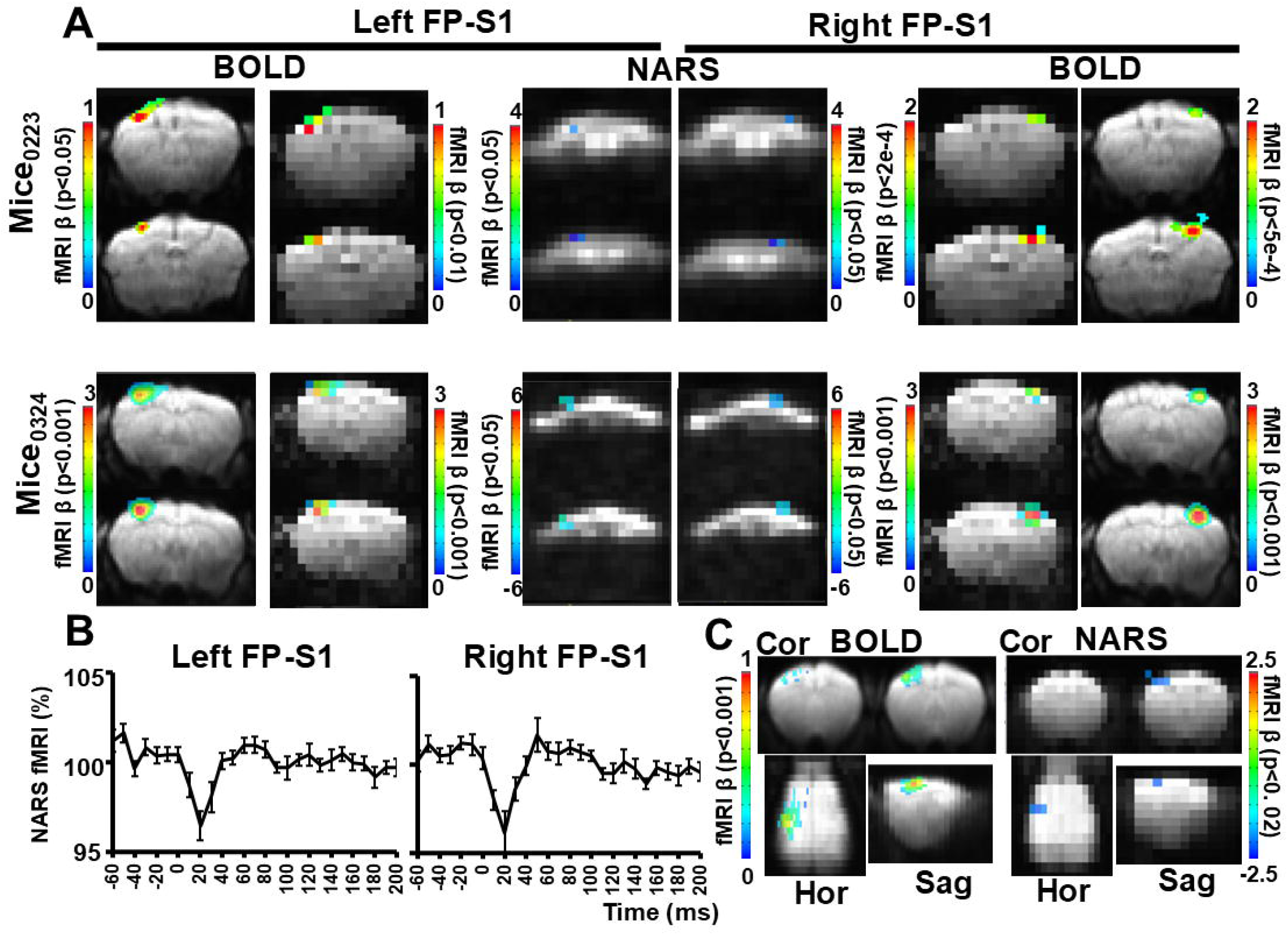
Reproducibility of NARS-fMRI responses in mice. (A). Bilateral NARS-fMRI activation of two representative mice. BOLD-fMRI activation map acquired in the same mice, demonstrating spatial correspondence with NARS-fMRI. Mouse0223 represents the FP-S1 activation with sequential bilateral forepaw stimulation with 100ms offset from the same trial, and mouse0324 represents FP-S1 activation of left and right forepaw stimulation at different trials. (B) Extracted FP-S1 NARS time course showing a rapid negative sodium response peaking within 10-30 ms from both hemispheres (mean±s.e.m, left FP-S1: n=11, right FP-S1: n=9). (C) Group-averaged NARS-fMRI responses across mice, confirming robust and reproducible millisecond-scale sodium dynamics (n=11).

To determine the temporal features underlying the NARS statistical maps, we fit the ultrafast series using a dynamic response function spanning varied onset times and durations. These models reproduce the NARS activation and yield the strongest correlations in FP-S1 for response peaks within ~10-30 ms of stimulus onset (Supp. Fig. 4 for rats; Supp. Fig. 5 for mice). Meanwhile, the stimulus onset was altered within the 300 ms acquisition window to test timing specificity from different animals. The evoked NARS decrease shifted in lock-step with the imposed onset, verifying that the millisecond-scale ^23^Na dynamics are time-locked to stimulation rather than sequence-related. (Supp. Fig. 6). Functional maps generated at multiple statistical thresholds for both BOLD and NARS-fMRI further demonstrate that the detected FP-S1 activation remains robust relative to background noise levels, reinforcing the reliability of the localized signal (Supp. Fig. 7). These combined rat and mouse data establish NARS-fMRI as a reliable, millisecond-resolution approach for mapping somatosensory cortex activation that is spatially concordant with BOLD yet temporally closer to neuronal events.

### Neuronal validation via simultaneous glutamate fiber photometry

To validate the neuronal origin of the NARS-fMRI signal, we performed simultaneous ^23^Na NARS-fMRI and glutamate (Glu) fiber-photometry recordings in FP-S1 using iGluSnFR in mice. iGluSnFR transients provide an optical proxy for fast synaptic activity, which can be used to track electrophysiological responses. In our preparation, spontaneous iGluSnFR events positively correlated with spontaneous LFP spikes, supporting the use of Glu photometry as a surrogate neural readout (Supp. Fig. 8A). Because the implanted optical fiber is paramagnetic, introducing little image distortion, it provides a favorable condition to be combined with rodent hemodynamic fMRI^68^. We observed robust evoked Glu signals together with well-defined BOLD-fMRI responses, indicating minimal electromagnetic interference and proper placement of the fiber to target the activated FP-S1(Supp. Fig. 8B).

Under repeated forepaw stimulation, trial-sorted Glu transients covaried with the amplitude of the NARS signal decrease across trials and animals. The larger evoked Glu responses consistently coincided with larger ^23^Na signal drops, demonstrating a tight coupling between synaptic glutamatergic activity and the fast NARS contrast (Fig. 4). Together with the spontaneous-state correlations above, these data indicate that elevated glutamatergic activity accompanies a greater engagement of fast-dephasing sodium microenvironments, consistent with an activity-dependent shift toward short-T_2_* pathways. In summary, concurrent iGluSnFR photometry and ^23^Na NARS-fMRI establish a stimulation-locked, glutamate-coupled NARS decrease in FP-S1, providing direct neuronal validation of the millisecond-scale NARS-fMRI contrast.

**Figure 4:**
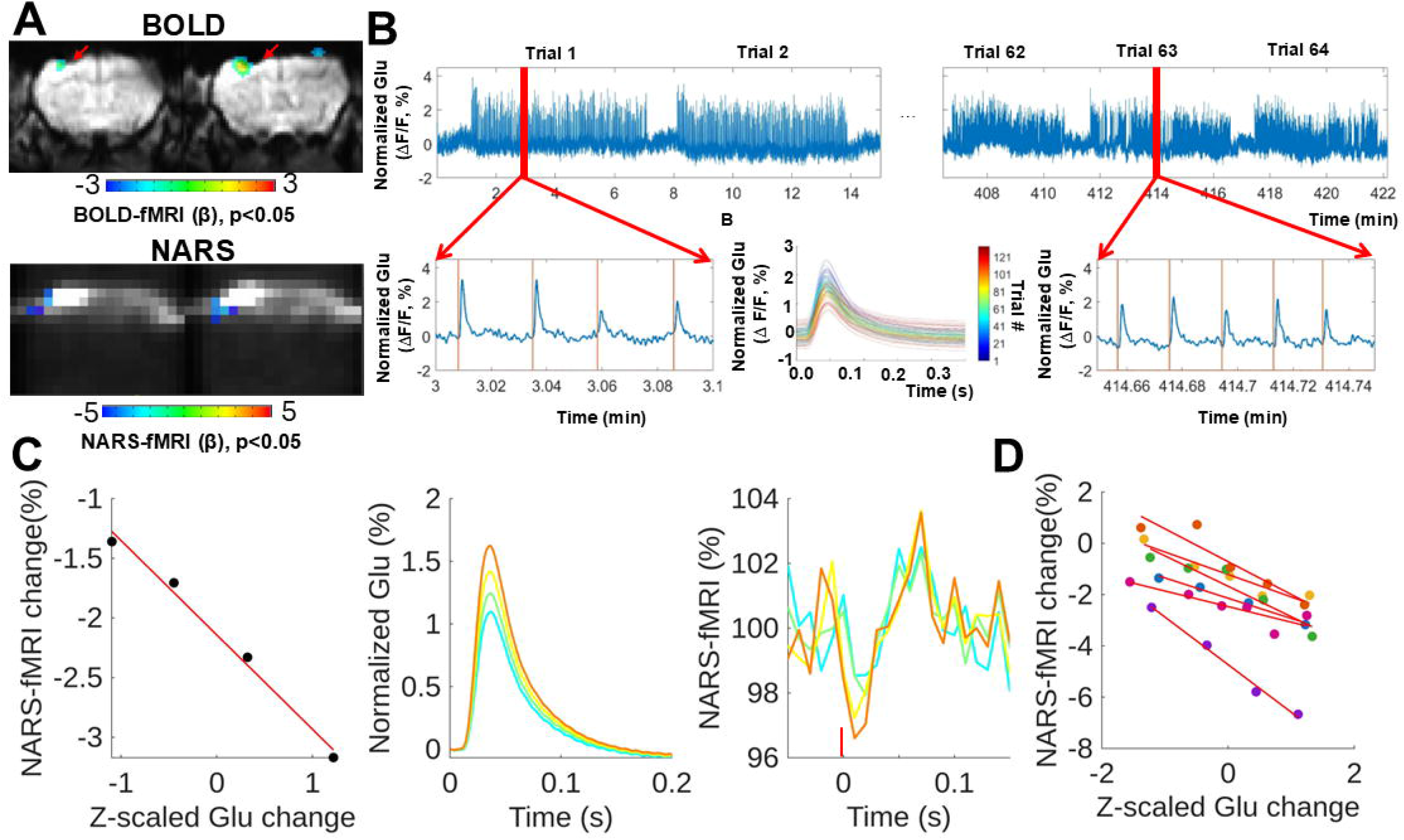
Neuronal validation of NARS-fMRI with simultaneous glutamate photometry. (A) BOLD and NARS fMRI functional maps of the FP-S1 with an optical fiber inserted (red arrow in the upper panel). The viral-injected area presented higher signal intensity in ^23^Na MRI images, suggesting the accumulation of sodium ions in the targeted brain regions. (B) Representative trace of the evoked Glu transients across different trials (#1…64). Enlarged images demonstrate the evoked Glu transient per stimulation pulse (inset shows the colored-coded average Glu responses per trial as a function of time). (C) Trial-by-trial scatter plot relating peak glutamate amplitude to peak NARS-fMRI signal change, demonstrating amplitude-dependent coupling. Trial-averaged glutamate (ΔF/F, %) and NARS-fMRI responses evoked by forepaw stimulation, showing temporally aligned glutamatergic transients and sodium signal decreases. (D) Linear regression of glutamate versus NARS-fMRI amplitudes across different animals (n=6), supporting a neuronal origin of the fast sodium response.

## Discussion

We introduce NARS-fMRI as a practical strategy for mapping neuronal-timescale activity in the rodent cortex. By integrating a figure-8 implantable RF coil with a reshuffled k-t 3D GRE acquisition at 14 T, we observed a robust negative ^23^Na response (~2-3%) in the forepaw somatosensory cortex, peaking 10-30 ms after stimulation. This signal spatially overlapped with BOLD-fMRI activation measured in the same animals and exhibited an amplitude-dependent negative correlation with simultaneously recorded glutamate transients, supporting a neuronal origin. Together, these findings establish NARS-fMRI as a mesoscale imaging modality that resolves fast neuronal dynamics and complements the hemodynamic contrast of BOLD-fMRI.

### Ultrafast fMRI and the pursuit of direct neuronal activity mapping

A central goal in functional neuroimaging is to move beyond indirect hemodynamic readouts and develop MRI-based contrasts that more directly reflect neuronal activity. A long-standing line of research has explored whether MRI could be used to directly sense electrophysiological activity^**69**^. One major approach is to use MRI to detect neuronal currents based on the transient magnetic fields capable of perturbing MR phase or frequency^70,71^. Quantitative modeling and phantom experiments have shown that the bio-magnetic fields produced by neural currents are on the order of nanotesla or smaller, corresponding to phase shifts lower than 0.2-0.4° ^72–75^. These estimates motivated further attempts to detect neuronal currents based on the phase shifts in turtle brains^76,77^, as well as feasibility studies targeting optic nerves^78^ or peripheral nerves^79^ in humans. A related line of work explored MRI signals arising from Lorentz force-driven tissue or ionic displacements^80,81^. In practice, these extremely small effects are readily overwhelmed by physiological noise, motion, and sequence-related artifacts, making robust and reproducible neural-current MRI challenging in vivo. As a result, sensitivity and artifacts fundamentally limit direct neural-current MRI.

Recent advances in ultrafast fMRI methods^35–37^ and multimodal validation frameworks provide a foundation for exploring non-hemodynamic MRI signals linked more closely to neuronal activity^13^. These developments, spanning early demonstrations of laminar-specific onset timing to later single-vessel and mesoscale implementations, have clarified the limits of vascular coupling and motivated renewed evaluation of non-hemodynamic MRI contrasts, including proton-based approaches to direct neuronal activity mapping^39,40^. In parallel, advances in implantable RF probe and resonator design have enabled millisecond-scale sampling of free-induction decay (FID) signals or single k-space line profiles by substantially boosting local SNR^47–50^. Combined with ultra-high magnetic fields, these technological advances remove key sensitivity barriers for *in vivo* ^23^Na MRI and make ultrafast sodium-based measurements feasible. This progress establishes NARS-fMRI as a complementary and experimentally tractable approach.

Instead of attempting to sense the extremely weak magnetic fields generated by neuronal currents, NARS-fMRI exploits the central role of sodium ions in action potentials and synaptic transmission, together with the strong sensitivity of ^23^Na MRI to microenvironment-dependent quadrupolar relaxation. The NARS response peaks on a 10-30 ms timescale and is temporally correlated with glutamatergic activity measured using iGluSnFR, supporting a neuronal origin. Spatially, NARS-fMRI localizes to regions defined by conventional BOLD mapping, yet exhibits a fundamentally different temporal profile, consistent with a non-hemodynamic contrast mechanism. At the voxel level, the NARS signal can be interpreted as the population-averaged superposition of many single-cell sodium transients. Although sodium influx at individual neurons occurs on sub-millisecond timescales, variability in spike timing, synaptic integration, and local diffusion across large neuronal populations produces a smooth, rapidly decaying mesoscale response. Together, these features place NARS-fMRI among ultrafast fMRI strategies that pursue neuronal specificity through biophysically grounded contrasts with direct links to ionic processes that can be independently validated.

### Biophysical origin of the NARS signal and modulation of single quantum T2*_short_

Most prior ^23^Na MRI studies in the brain have focused on measuring tissue sodium concentration changes associated with pathology, including ischemia^82–84^, brain tumors^85–87^, and neurodegenerative disorders^88–92^. In these contexts, sodium MRI has been used to detect disruptions of ionic homeostasis arising from impaired membrane integrity, energy failure, or altered cellular and extracellular volume fractions. While these approaches are powerful for characterizing disease-related sodium accumulation, they primarily probe slow processes evolving over minutes to hours^28,93,94^. By contrast, NARS-fMRI targets rapid, activity-dependent sodium dynamics and does not require large alterations in bulk tissue sodium concentration.

A central observation of this study is a robust 2-3% negative NARS signal change detected at ultrashort TE (~1 ms). Importantly, this effect cannot be explained by bulk intracellular sodium accumulation. Electrophysiological studies have shown that the net intracellular sodium concentration change associated with a single action potential is extremely small, typically well below 0.1% of baseline intracellular sodium when averaged over the cell volume, even during repetitive firing^18,95,96^. At the voxel scale, such changes are orders of magnitude too small to account for the observed NARS signal amplitude. Instead, the data support a mechanism in which neuronal activation induces a transient redistribution of sodium ions among microenvironments with distinct relaxation properties^97^. During an action potential, sodium ions rapidly enter neurons through voltage-gated channels, producing highly localized and short-lived increases in sodium density near membranes, ion channels, transporters, and cytoskeletal or protein-rich regions^98,99^. In parallel, sodium is transiently depleted from the perisynaptic extracellular space and redistributed along highly confined extracellular microdomains. Although individual sodium-macromolecule interactions are brief, the concerted effect of synchronized transmembrane sodium flux and perisynaptic redistribution shifts a fraction of sodium ions toward more restricted or partially ordered environments on both sides of the membrane^100^.

Such microenvironments enhance quadrupolar relaxation through increased local electric field gradients and altered correlation times, thereby increasing the relative contribution of fast-relaxing sodium components. At ultrashort echo times, even modest changes in the fractional occupancy of these fast-relaxing pools can lead to measurable signal reductions, providing a biophysically plausible explanation for the observed shortening of T2*short and the negative NARS response. In this framework, diffusion primarily shapes the spatial and temporal smoothing of the voxel-level signal but does not constitute the dominant contrast-generating mechanism.

This interpretation is consistent with, and supported by, a growing body of cellular sodium MRI studies that move beyond bulk tissue sodium concentration measurements. Earlier efforts to separate intracellular and extracellular sodium pools using shift reagents^27,101,102^ or multiple-quantum filtering^102–104^ provided important insights but were limited by sensitivity, specificity, or *in vivo* applicability. More recently, established relaxation-exchange spectroscopy approaches combined with multisite exchange modeling have been applied to living cells to quantify transmembrane sodium transport rates and intracellular relaxation parameters using endogenous ^23^Na relaxation contrasts^105^. These studies demonstrate that physiologically relevant sodium dynamics, including sodium influx and exchange across the membrane, can be inferred from relaxation-weighted measurements without exogenous agents. Together, these results suggest a clear mechanistic pathway for future NARS-fMRI studies to integrate cellular-scale sodium transport models with *in vivo* ultrafast sodium imaging, enabling more quantitative links between neuronal activity, transmembrane sodium flux, and relaxation-based MRI signals.

### Technical considerations: physiological noise, sequence artifacts, and RF excitation

Ultrafast fMRI places stringent demands on artifact control, as physiological fluctuations that are negligible in conventional fMRI can dominate ^23^Na-based fMRI signals. Respiration is a particularly critical confound for sodium imaging, as chest and neck motion induce global B_0_ perturbations that can alias into the reconstructed time series, especially when k-space data are reshuffled across time. In addition, respiration-induced changes in RF coil loading can introduce apparent signal fluctuations. In this work, respiration-gating synchronized to the stimulation paradigm was essential for suppressing these confounds. Combined with an implanted sodium coil embedded in dental cement, this approach minimized both B_0_ and B_1_ instabilities. The robustness of the NARS response to stimulus-onset times further supports a time-locked neuronal origin rather than a periodic physiological or sequence artifact.

Potential sequence-related confounds become more consequential at ultrashort TR and TE, even for ^23^Na 3/2 spins, where the signal is dominated by non-equilibrium magnetization and by how residual transverse pathways are dephased or inadvertently refocused across rapid RF and gradient trains^106^. In this regime, small imperfections in spoiling, gradient timing, or local off-resonance can introduce structured signal fluctuations unrelated to neuronal activity. Recent work has further emphasized that ultrafast MRI signals may contain steady-state-dependent oscillatory components that reflect frequency offsets rather than true stimulus-evoked dynamics^67^ The consistency of the observed NARS response across repeated trials with highly varied respiration-gated inter-epoch intervals (Supp Fig 2) and timing manipulations (Supp Fig 6) argues against such effects as the primary source of the signal, but future studies incorporating additional controls, such as alternative k-space trajectories or multi-echo acquisitions, will further strengthen mechanistic interpretation.

With respect to RF excitation, the 12-20 kHz bandwidths used here are far larger than any physiologically relevant ^23^Na quadrupolar splitting observed *in vivo*, which typically fall in the tens-to-hundreds of hertz range^107^, and at most reach a few kilohertz only in highly ordered tissues such as cartilage^108^. As a result, the entire effective ^23^Na spectral envelope, including the center transition and any broadened satellite components, is fully encompassed by the applied RF pulses. Although residual interactions in partially ordered environments may modestly modulate relaxation behavior, these effects are not spectrally resolved *in vivo* and manifest solely as relaxation-domain phenomena rather than frequency-selective components. Consequently, 3/2 spin-related quadrupolar splitting does not pose any risk of differential excitation and is not a confounding factor for NARS fMRI.

### Limitations and prospects for translation to humans

The reshuffled k-t space acquisition enabled ultrafast sampling of repetitively stimulated brain regions. This paradigm emphasizes neuronal responses that are tightly time-locked to stimulation and limits sensitivity to activity that is weakly driven or rapidly desensitized. Instead, such non-phase-locked activity would otherwise act as physiological noise when mapping NARS-fMRI signals. In addition, ultra-fast acquisitions are inherently sensitive to motion and temporally aliased artifacts and may capture non-physiological oscillations if not carefully controlled. Respiration-gating and systematic variation of stimulation paradigms across species are therefore essential to establish robustness and reproducibility. At present, NARS-fMRI relies on ultra-high magnetic field strength and specialized RF hardware to achieve sufficient SNR at ultrashort echo times. An important next step is to integrate line-scanning fMRI^37^ to extract real-time NARS-fMRI fluctuations with improved temporal specificity. Combining this approach with simultaneous neuronal recordings and implantable, wirelessly amplified micro-coils will further boost local sensitivity and enable direct cross-modal validation^47–50^. Although a microenvironment-based mechanism provides a plausible interpretation of the NARS signal, it remains a working hypothesis and requires further validation. Disentangling neuronal sodium dynamics from vascular or metabolic effects requires experimental systems with reduced hemodynamic confounds. Cerebral organoids provide such a platform. They preserve neuronal and synaptic activity while lacking mature vasculature, enabling isolation of activity-dependent sodium redistribution at the cellular and network levels. Integration of NARS-fMRI with electrophysiology, optical reporters, and cellular sodium modeling in organoids will therefore be critical for mechanistic validation. Despite these challenges, the translational potential of NARS-fMRI is compelling. Human implementation at high field (e.g., 7T), combined with optimized sodium receive arrays and ultrashort-TE trajectories, could enable mesoscale mapping of neuronal activity with temporal specificity beyond that of conventional BOLD fMRI.

## Methods

### Animals

Adult female C57BL/6 mice (3-5 months old; n = 29) and female Long-Evans rats (4 weeks old; n = 14) were used in this study. The exact number of animals included in each experiment is provided in the corresponding figure legends. Some animals contributed to more than one experiment (e.g., T_1_ or T_2_* mapping and NARS-fMRI), and this overlap is indicated where relevant. Animals used exclusively for RF-coil optimization or viral-vector validation were not included in the reported totals, as these procedures were either part of previously published work or performed in parallel studies. For transparency and reproducibility, representative datasets are labeled by acquisition date to facilitate raw-data retrieval. Animals were group-housed (3-4 per cage) under a 12-h light/dark cycle with ad libitum access to food and water. All procedures were approved by the Massachusetts General Hospital Institutional Animal Care and Use Committee (IACUC) and complied with the National Research Council’s *Guide for the Care and Use of Laboratory Animals*.

### Animal Surgery

#### RF coil implantation (mice and rats)

Mice and rats underwent surgery to affix a custom-built RF coil to the skull. Anesthesia was induced with 5% isoflurane delivered in medical air supplemented with O_2_ (15-20%) for 2– 3 min, and maintained at 1.5-2.0% isoflurane with respiratory rate used to monitor anesthetic depth. Ophthalmic ointment was applied to protect the corneas, and body temperature was maintained at 37 °C using a thermostatically controlled warming pad throughout the procedure. Animals were placed in a stereotaxic frame and secured with ear bars and a bite bar to ensure stable head positioning. Ethiqa XR (buprenorphine extended-release), 3.25 mg kg^−1^, s.c., was administered once pre-operatively; per protocol, no additional post-operative analgesia was required due to its prolonged effect.

The scalp was shaved and sterilized with alternating ethanol and iodine swabs. A midline incision was made to expose the skull over the region corresponding to the RF-coil footprint. Residual soft tissue was removed and the skull surface was cleaned sequentially with 0.3% H_2_O_2_ and PBS, then allowed to dry. The RF coil was positioned directly over the skull above the targeted brain region, and its tuning was verified at the ^23^Na Larmor frequency at 14 T. The coil was temporarily held in place while a thin layer of cyanoacrylate adhesive was applied to bond it to the skull; curing required approximately 5–8 min. Two-part dental cement (Stoelting Co., Wood Dale, IL) was then mixed and applied to encapsulate the coil and exposed bone, taking care to secure the base of the coil firmly and prevent cement from flowing toward the eyes. After cement placement, the scalp edges were approximated and closed using tissue adhesive. For mice, once the cement had fully hardened (~10 min), animals were removed from the stereotaxic frame and transferred to a warmed recovery cage until fully ambulatory. Mice were then returned to their home cages and allowed to recover for at least one week to ensure adequate neck-muscle strengthening and stable head posture before imaging. If animals experience head implant removal, infection, etc. they will be euthanized based on the approval procedure from the protocol. In rats, RF-coil implantation was performed acutely on the same day as the terminal fMRI experiment. After coil fixation and stabilization, rats remained under anesthesia and were immediately transported for imaging.

#### Viral vector injection (mice only)

Viral injections were performed 5 weeks prior to RF-coil and optical-fiber implantation to allow sufficient transgene expression. Mice were anesthetized with isoflurane (induction at 5% in medical air with supplemental 15-20% O_2_; maintenance at 1.5-2.0% adjusted to respiration). Ethiqa XR (3.25 mg kg^−1^, s.c., was administered once pre-operatively). Ophthalmic ointment was applied to prevent corneal drying, and body temperature was maintained at 37 °C on a thermostatically controlled warming pad. Animals were positioned in a stereotaxic frame (ear bars and bite bar), and the skull was leveled by aligning bregma and lambda in the dorsoventral and mediolateral planes.

The scalp was shaved and disinfected with alternating ethanol and iodine swabs. A midline incision exposed the skull, and stereotaxic coordinates were marked relative to bregma (target region: AP, −0.5mm, ML, ±2.3mm) to target FP-S1. A small burr hole was drilled, and a 33-gauge needle coupled to a nano-injector with 10µl microsyringe was loaded with the viral vector (AAV9-hSyn.iGluSnFr.WPRE.SV40, Addgene, 98929-AAV9). The injector was lowered to 0.6mm, and 200nL was delivered at 100 nL/min. For 3-stop placements, aliquots were deposited while retracting in 200 µm steps to minimize reflux; the injector remained in place for 5 min after each deposit before slow withdrawal. Burr holes were sealed with sterile bone wax, and the incision was closed with interrupted sutures. Mice were monitored on a warmed surface until recovery and then returned to home cages. Post-operative monitoring was performed at least daily for 72 h.

Implantation surgery for imaging (RF coil and optical fiber) was scheduled no earlier than 5 weeks post-injection to ensure robust, stable expression. After positioning the RF coil above the skull following the surgical procedure as mentioned above, the optical fiber (Thorlabs, low-OH, 200µm diameter) was slowly inserted through the small burr hole drilled based on the prior coordinates to target the FP-S1 at 400-500µm depth. And, the same fixation procedure will be applied to secure both the RF coil and optical fiber above the mouse skull. The multi-fMRI procedure will be performed on the same day, followed by ex vivo histology to verify the Glu expression after brain perfusion using the procedure described previously^68^.

### Animal preparation for fMRI

For NARS/BOLD-fMRI experiments, mice and rats were imaged under free-breathing medetomidine anesthesia in combination with low-dose isoflurane. Animals were initially anesthetized with isoflurane (5% induction in medical air with supplemental 15-20% O_2_). A subcutaneous bolus of medetomidine (0.05 mg kg^−1^) was then administered, after which isoflurane was reduced and maintained at 0.5-1.0%, titrated slowly over the course of the imaging session (6-10 h total duration) to maintain physiological stability. Anesthesia was maintained with a continuous low-dose medetomidine infusion (~0.1 mg kg^−1^ h^−1^, s.c.). Ophthalmic ointment was applied to protect the cornea, and body temperature was maintained at 37 °C with a thermostatically regulated warm-air heating system.

Animals were positioned in a custom MRI-compatible head holder incorporating ear bars and a bite bar to minimize motion while preserving natural respiration. No paralytics were used. Physiological parameters, including respiration rate and body temperature, were continuously monitored using a small-animal monitoring system throughout the procedure (SA Instruments, NY). Once stable under medetomidine/isoflurane anesthesia, animals were transported into the MRI bore for MR scanning. For mice, simultaneous fiber-photometry recordings were performed during fMRI sessions. Photometry signals were time-synchronized with fMRI acquisition to enable integrated NARS-fMRI measurements.

### Electrophysiological and glutamate photometry recording

Electrophysiological recording and glutamate fiber photometry were performed following stereotaxic implantation of a 200 µm low-OH optical fiber (Thorlabs, 200EMT) and a tungsten microelectrode (1 MΩ, ~100 µm; FHC). Implants were targeted to deep layers of the forepaw primary somatosensory cortex (FP-S1) using species-specific coordinates (rats: caudal 0.2 mm, lateral 4.0 mm, ventral 1-1.2 mm; mice: caudal 0.5 mm, lateral 2.3 mm, ventral 0.6 mm). For simultaneous recordings, the dura was carefully removed, and the electrode was affixed adjacent to the optical fiber such that the electrode tip and fiber tip sampled the same local microcircuit. Reference and ground leads were connected to a skull screw placed over the right cerebellum. Both evoked and spontaneous local field potentials (LFPs) were recorded using a Biopac EEG module (gain 5,000; band-pass 0.02-100 Hz; sampling rate 5 kHz). Glutamate photometry signals derived from expressed iGluSnFR sensors were acquired using a custom-built fiber-photometry system^68^ and digitized through the analogue input module of the Biopac MP-150 interface. The Biopac system also recorded synchronized TTL pulses to ensure temporal alignment of fluorescence acquisition, electrophysiology, and external stimulation. Forepaw electrical stimulation (300 µs biphasic pulses at 3 Hz for 4 s or 30 s, 1.5-2.5mA) was delivered through interdigital needle electrodes using a stimulus isolator (WPI A365), with timing controlled by a Master-9 (A.M.P.I). Throughout the recording sessions, body temperature and respiratory rate were continuously monitored using a small-animal physiological monitoring system. Custom MATLAB scripts were used for offline extraction of glutamate transients, LFP responses, and stimulation timestamps, enabling quantitative analysis of both spontaneous and evoked signal dynamics.

### Immunohistochemistry

After imaging or electrophysiological/photometry recording, mice were transcardially perfused with ice-cold PBS followed by 4% paraformaldehyde (PFA). Brains were post-fixed overnight in 4% PFA at 4 °C and then cryoprotected in 15% and subsequently 30% sucrose in PBS until fully equilibrated. Tissue was sectioned at 30 µm on a cryostat (Leica CM3050S). Floating sections were rinsed in PBS and mounted directly without antibody incubation. DAPI-containing mounting medium (VectaShield, Vector Laboratories) was applied to label nuclei and cover the sections. Glu-sensor fluorescence (iGluSnFR) was visualized directly from the endogenous reporter signal without immunoenhancement. Images were acquired using a wide-field fluorescence microscope to assess overall sensor expression and fiber-implant location. High-resolution confocal images (Leica SP2, 10× objective) were collected when needed to confirm local expression in the targeted FP-S1 region.

### MRI methods

#### MRI systems

All experiments were performed on horizontal bore MRI scanners (Magnex Scientific, UK, 130mm bore size) at the Athinoula A. Martinos Center for Biomedical Imaging (Charlestown, MA, USA). The 14 T (600 MHz for ^1^H, ~158MHz for ^23^Na) system was equipped with a Bruker Avance NEO console running ParaVision 360 v3.3 and a micro-imaging gradient set (Resonance Research, Inc.) providing peak gradient strength 1.2 T m^−1^ over a 60-mm diameter.

#### Home-built implantable RF coils (^1^H and ^23^Na)

Custom implantable RF coils were fabricated in-house on thin PCB substrates following a previously described PCB-based framework^55^. Both coils shared identical construction and matching approaches, differing only in loop geometry and resonance. For ^1^H imaging, a single-loop circular transceiver (≈ 20 mm diameter) was tuned and matched to 50 Ω at the ^1^H Larmor frequency at 14 T (~600 MHz). For ^23^Na imaging, a compact figure-8 transceiver comprising two adjacent circular loops (4-5 mm diameter each) wired in series opposition was tuned to the ^23^Na Larmor frequency at 14 T (~158 MHz). Resonance and match were set using distributed NP0/C0G capacitors on the PCB to adjust the effective loop inductance and capacitance. Both coils were connected to a miniature 50 Ω coax with integrated strain relief in the PCB layout. When mounted above the head, the B_1_ fields of the single-loop ^1^H and figure-8 ^23^Na coils are approximately orthogonal, so no additional decoupling was required in practice.

##### MRI data acquisition

i). For mouse studies, the initial anatomical scan used a multi-slice 2D FLASH sequence: TE = 2.72 ms, TR = 500 ms, NA = 2, matrix = 120×120 over a 12×12 mm^2^ FOV, 24 slices at 0.5 mm thickness, providing the structural reference aligned to the subsequent functional scans. Multi-slice 2D gradient-echo EPI for BOLD fMRI with TE/TR = 5.7 ms/500 ms, two-shot segmentation, acquisition bandwidth 151,515 Hz, 60×60 matrix over a 12×12 mm^2^ FOV (200 µm in-plane), 24 axial slices at 0.5 mm thickness, and 410 repetitions for functional time-series. The ^23^Na-based NARS fMRI acquired in the same orientation but with slightly different geometry (FOV = 12×9×12 mm^3^, matrix = 20×15×15 or 20×15×10; voxel size 0.6×0.6×0.8 mm^3^ or 0.6×0.6×1.2 mm^3^), using TR/TE = 10 ms/1.005 ms, RF excitation pulse duration 0.25 ms (16.8 kHz RF bandwidth), 1.3 ms acquisition window with a 10 kHz readout bandwidth, and the same movie-loop framework (30 frames).

In addition, ^23^Na T1 and T2* mapping was performed using a 3D gradient-echo MRI sequence. The same acquisition protocol was repeated multiple times with different TR or TE to sample the sodium signal decay. For T2* mapping, different TEs were used from 0.8ms to 80ms with following parameters: TR, 100 ms, RF pulse duration, 0.25 ms, bandwidth, 16.8 kHz; flip angle 90°, NA, 100; FOV: 12 × 12 × 12 mm^3^, matrix size: 15 × 15 × 10, corresponding to a spatial resolution of 0.8 × 0.8 × 1.2 mm^3^. For T1 mapping, TE was fixed at 1 ms, and a series of scans was acquired with TR varied from approximately 5 ms to 350 ms. All other parameters were identical to those used for T2* mapping.

ii). For rat studies, the anatomical reference used a multi-slice 2D FLASH sequence (TE/TR = 3.32 ms/500 ms, NA = 1, matrix 192×192 over a 19.2×19.2 mm^2^ FOV, 12 slices at 0.6 mm thickness; in-plane resolution 100 µm) acquired with interlaced slice order. BOLD fMRI was acquired with a multi-slice 2D gradient-echo EPI sequence (TE/TR = 5.4 ms/500 ms, 2-shot segmentation, acquisition bandwidth 151,515 Hz, matrix 64×64 over a 19.2×19.2 mm^2^ FOV yielding 300 µm × 300 µm in-plane resolution, 12 axial slices at 0.6 mm thickness, 410 repetitions). ^23^Na-based NARS fMRI was then acquired in the same orientation but with a slightly different geometry: FOV 15×9×7.2 mm^3^, matrix 25×15×6 (encoded 18×15×6) for 0.6×0.6×1.2 mm^3^ voxels, using TR/TE = 10 ms/1.25 ms, RF excitation pulse duration = 0.2 ms with 21 kHz RF bandwidth, acquisition window = 1.98 ms, spectral/readout bandwidth = 9.09 kHz, and a movie-loop of 60-20 frames.

### Stimulation paradigm and respiration gating

#### BOLD fMRI

Forepaw electrical stimulation (biphasic pulses, 300 µs, 1.5-2.0 mA) was delivered at 3 Hz for 8 s per block. The image-embedded block schedule for BOLD used 10 baseline scans, 1 stimulation-trigger scan, and 39 rest scans, repeated for 10 epochs (total 410 TRs). No respiration gating was used for BOLD. 3-8 trials were acquired across different animals.

#### NARS-fMRI (^23^Na)

Stimulation was respiration-gated and delivered as one electrical pulse per respiratory cycle, enabling the sequence trigger module (per-phase-step) for physiological synchronization. To probe timing sensitivity, stimulation onset within each movie loop varied between 100 ms, 150 ms, and 200 ms after loop start (Supp Fig. 6). In mice, sodium runs followed the method’s loop framework of 30 frames per loop based on the respiratory rate (120-160 bpm), lower than 3Hz (Supp Fig 2). In rats, given lower respiratory rates (40-60bpm), we used multiple short epochs each comprising 30 scans per epoch with one pulse per epoch, while recording 60/120-frame movie loops to capture respiration-locked sodium dynamics. We acquired 90-300 trials across animals.

### Multi-modality data analysis

All analyses were performed on Linux using AFNI, MATLAB, and custom Bash scripts. BOLD and NARS-fMRI data were processed in parallel pipelines that shared a FLASH-based anatomical registration framework to enable accurate cross-modality alignment and overlay. Acquisition-specific features, including respiration-gated sodium movie-loop acquisitions, stimulus-onset jitter, and reconstruction or averaging schemes, were preserved throughout the processing workflow. Electrophysiological and fiber photometry data were processed using custom MATLAB scripts, with multimodal synchronization achieved via hardware trigger outputs generated by the EPI and ultrafast ^23^Na GRE fMRI sequences.

#### SNR estimation for 23Na MRI datasets

SNR for NARS-fMRI was estimated directly from raw magnitude images acquired at fixed TE for NARS fMRI or with variable TRs and TEs for T_2_* and T1 mapping. For each acquisition, signal intensity was measured within an anatomically defined region of interest (ROI), while noise was estimated from a background region outside the sample that was free of visible signal and artifacts. SNR was computed as

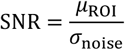

where *μ*_ROI_ denotes the mean signal intensity within the ROI and *σ*_moise_ denotes the standard deviation of the background noise. For magnitude images, noise estimates were corrected for the Rician bias.

#### Functional ^1^H/^23^Na MRI data analysis

For BOLD fMRI analysis, a thorough data analysis pipeline is described previously^54^. Briefly, the EPI time series were converted to AFNI format and rigidly aligned with 3dvolreg function and registered to the FLASH-based anatomical template with the 3dAllineat function. Spatial smoothing was avoided (beyond minimal interpolation during alignment) to maintain mesoscale specificity, in line with ultra-high-field rodent fMRI practices^109^. For direct comparison with sodium maps, BOLD images and statistics were additionally resampled to the sodium (NARS) grid so all overlays and ROI measurements were performed at matched voxel size. The general linear modeling was performed with the 3dDeconvolve function using compact BLOCK basis. Subject t-threshold-based β-maps were propagated to template space for group inference and cross-modality overlays, including versions on the NARS grid for voxel-parity checks.

For NARS fMRI analysis, sodium (^23^Na) movie-loop series were reconstructed and smoothed (0.6-0.8 mm blur size) to stabilize SNR without compromising mesoscale localization. A baseline was computed from frames 3–8 of each loop, framewise data were converted to percent signal changes, and the first three frames were discarded to remove steady-state transients. Since the 23Na MRI image datasets are acquired with the same geometry as EPI datasets, for group analysis, we applied the same transformation matrix to align all ^23^Na MRI datasets to the anatomical template. To enable GLM modeling within AFNI’s seconds-based framework, the temporal sampling of the ^23^Na time series was adjusted to be compatible with AFNI’s built-in GLM analysis while preserving the native temporal dynamics of the sodium signal. The stimulation profile was aligned to this adjusted timeline, and BLOCK-based regressors were used for each stimulus channel, capturing the brief sodium responses and yielding robust voxelwise β/t estimates for group inference and overlays. To validate the millisecond-scale timing of the NARS-fMRI response independently of the AFNI GLM, we additionally performed a native-time sodium response analysis using gamma variate–like ideal functions in MATLAB. Gamma variate curves are defined as:

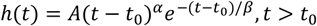

(A is amplitude, *t*_O_ is the onset delay, *α* controls rising slope, and *β* controls width).

which yield smooth unimodal responses with adjustable onset and width. In practice, we generated families of brief gamma-like kernels with 10, 15, and 20 ms durations and applied onset shifts of 0–20 ms. Each kernel was normalized and correlated with the sodium ROI time courses sampled at the true 10 ms movie-loop rate, with the first three frames discarded to remove unsaturated in-flow transients. Across animals, the best GAM fit closely matched the AFNI BLOCK solution in timing and shape (Supp Fig 4,5), supporting the pseudo-BLOCK approach for GLM while preserving millisecond specificity via native-time analysis.

For both modalities, subject-level β/t maps were transformed to template space and entered one-sample t-tests (AFNI 3dttest++) for group inference. Group overlays are presented at voxelwise p < 0.05, p < 0.01, and p < 0.001 (two-tailed), as labeled in figure panels. Resampling-based stability checks for sodium (random subsets of runs with epoch averaging and refitting over 64 repetitions) reproduced the deterministic GLM patterns.

#### Amplitude-dependent Glu with simultaneous LFP and NARS-fMRI analysis

For spontaneous Glu-LFP coupling analysis, Glu fluorescence transients and LFP signals were recorded simultaneously. Glu traces were baseline-normalized and detrended, and spontaneous Glu transients were identified using a threshold-based detection criterion. For each detected event, the peak Glu amplitude was extracted within a fixed window (~100ms). Corresponding LFP segments were aligned to the same event times, and LFP response amplitude was quantified as the peak-to-trough value within a predefined peri-event window. Event-wise peak amplitudes from the two modalities were then compared using scatter plots, and their relationship was quantified using linear regression. This analysis establishes a direct correspondence between spontaneous Glu signaling and underlying electrophysiological activity.

For simultaneous Glu photometry and NARS-fMRI experiments, a scalar glutamate response amplitude was computed as the peak ΔF/F percent change relative to a pre-stimulus baseline within a fixed post-stimulus window for each trial. In parallel, NARS-fMRI signals were extracted from activated FP-S1 and quantified as the percent signal change relative to a baseline period. To assess amplitude dependence, trials were ordered by glutamate response amplitude and grouped into consecutive bins by averaging neighboring trials. For each bin, mean glutamate and mean NARS-fMRI response amplitudes were calculated. The relationship between glutamate and NARS-fMRI amplitudes was then evaluated using linear regression. To account for inter-animal variability in iGluSnFR expression, fiber placement, and optical transmission efficiency, z-score normalization was applied to glutamate amplitudes before regression. This approach reduces trial-to-trial noise while preserving systematic amplitude scaling between modalities. Across analyses, linear fitting was used solely to assess amplitude-dependent relationships between Glu and LFP, or Glu and NARS signals, providing complementary validation that NARS-fMRI signal magnitude scales with neuronal and synaptic activity across electrophysiological, optical, and MRI readouts.

#### T1 mapping

Multi-TR gradient-echo datasets were processed in MATLAB to estimate T_1_ from SNR-normalized recovery curves from cortical ROIs. T_1_ was obtained by non-linear least squares to a mono-exponential recovery model,

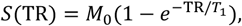

where *M*_0_is the equilibrium magnetization and *T*_1_is the longitudinal relaxation time. Fits used bounded Levenberg-Marquardt optimization with data-driven initial conditions, and *T*_1_was taken directly from the fitted exponential time constant. For each dataset, fitted and measured recovery curves were generated, and the resulting cortical T_1_ values were produced for group analysis.

#### T2 mapping

^23^Na gradient-echo datasets were processed in MATLAB to estimate T_2_* from SNR-normalized decay curves extracted from cortical ROIs. Sodium signal decay was modeled using a biexponential quadrupolar relaxation model of the form:

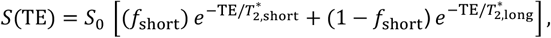

where 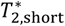 and 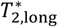 represent the fast and slow transverse relaxation components associated with ^23^Na quadrupolar dephasing, and *f*_short_is the fractional amplitude of the short-T_2_* component.

Two fitting strategies were applied. In the fixed-fraction model, the short-component weight was constrained to 0.6 to stabilize estimation for the spin 3/2 ^23^Na signal under quadrupolar relaxation in the fast-motion regime, where the central and satellite transitions contribute in an approximately 0.6:0.4 ratio^23,24^. This constraint provides a physically motivated prior that stabilizes bi-exponential fitting in vivo, particularly under limited SNR and echo sampling, while preserving the standard fast/slow T_2_* decomposition. In the unconstrained model, the fractional contributions of the short- and long-T_2_* components were treated as free parameters during optimization, together with 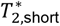 and 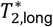. This approach allows the fitting to capture tissue- and voxel-dependent deviations from the idealized quadrupolar model arising from microenvironmental heterogeneity. Free-fraction bi-exponential fitting has been widely adopted in prior ^23^Na MRI relaxometry studies of brain and other tissues and was included here to assess the robustness of relaxation estimates to modeling assumptions^22,65,110,111^. In both cases, bounded Levenberg–Marquardt nonlinear least-squares fitting was used with data-driven initialization, yielding estimates of 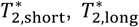, and the corresponding component amplitudes. For each dataset, fitted curves were overlaid on measured decay points as quality control, and cortical T_2_*short, T_2_*long, and component fractions were summarized as mean ± SEM for group analysis.

## Author Contributions

X.Y. built the concept, acquired and analyzed data, wrote the paper; X.L. analyzed data, G. Y. built RF coils and acquired data, Y.J. acquired ephys data, N.P. performed animal experiments, D.H. provided technological support, XA. Z. acquired data and performed animal experiments.

### Acknowledgements

This grant is supported by NIH grants (RF1NS124778, R01NS122904), NSF grant 2123971, and the S10 instrument grants (S10 MH124733, S10OD036211) to the Martinos Center. We thank Sascha Koehler and Rolf Pohmann for pulse sequence modification, thank …

