## supplementary figure 1-7 and legends for "Neuronal-Activity-Related Sodium (NARS) fMRI Reveals Millisecond Neuronal Dynamics Beyond Hemodynamic Readouts"

Running title: NARS-fMRI resolves millisecond neuronal dynamics

Xin Yu*, Xiaochen Liu, Grace Yu, Yuanyuan Jiang, Nivetha Pasupathy, David Hike, Xiaoqing Alice Zhou

### Supplementary Fig 1

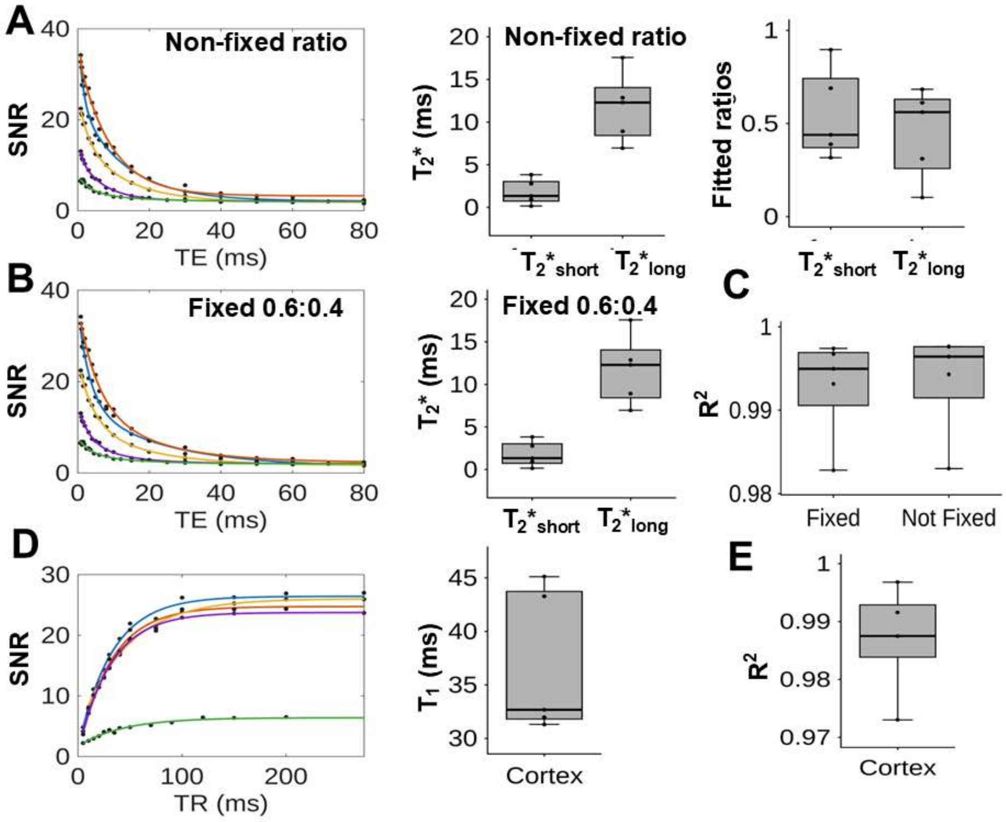


**Supplementary Fig 1. *In vivo* T1 and T2* mapping of the mouse cortex with 14T.**

(A) Representative cortical $^{23}$Na $T_2$* decay curves fitted with a bi-exponential model allowing component fractions to vary freely. Free-fraction fitting yielded $T_2{*}_{short}$ = 1.80 ± 0.66 ms, $T_2{*}_{long}$ = 11.7 ± 1.82 ms, with mean component weights of $f_{short}$ = 0.45 ± 0.11 and $f_{long}$ = 0.55 ± 0.11 (mean ± s.d., n=5), reflecting tissue-dependent heterogeneity and partial-volume effects.

(B). Representative cortical $^{23}$Na $T_2$* decay curve fitted with a bi-exponential model using a fixed fast-to-slow amplitude ratio of 0.6:0.4. The fixed-ratio fits yielded $T_2{*}_{short}$ = 3.91 ± 0.41 ms and $T_2{*}_{long}$ = 16.7± 1.94 ms (mean ± s.d., n=5), consistent with quadrupolar relaxation of spin-3/2 sodium in biological tissue.

(C) Both fitting models show $R^2$ higher than 0.99. (D/E) Representative inversion-recovery curves acquired from cortical ROI with mono-exponential fits. The mean cortical $T_1$ was 36.9 ± 3.01ms (mean ± s.d., n=5), with high goodness-of-fit ($R^2$ = 0.99 ± 0.01), indicating reliable $T_1$ estimation under ultrashort-TE acquisition conditions.

**Supplementary Fig 2**

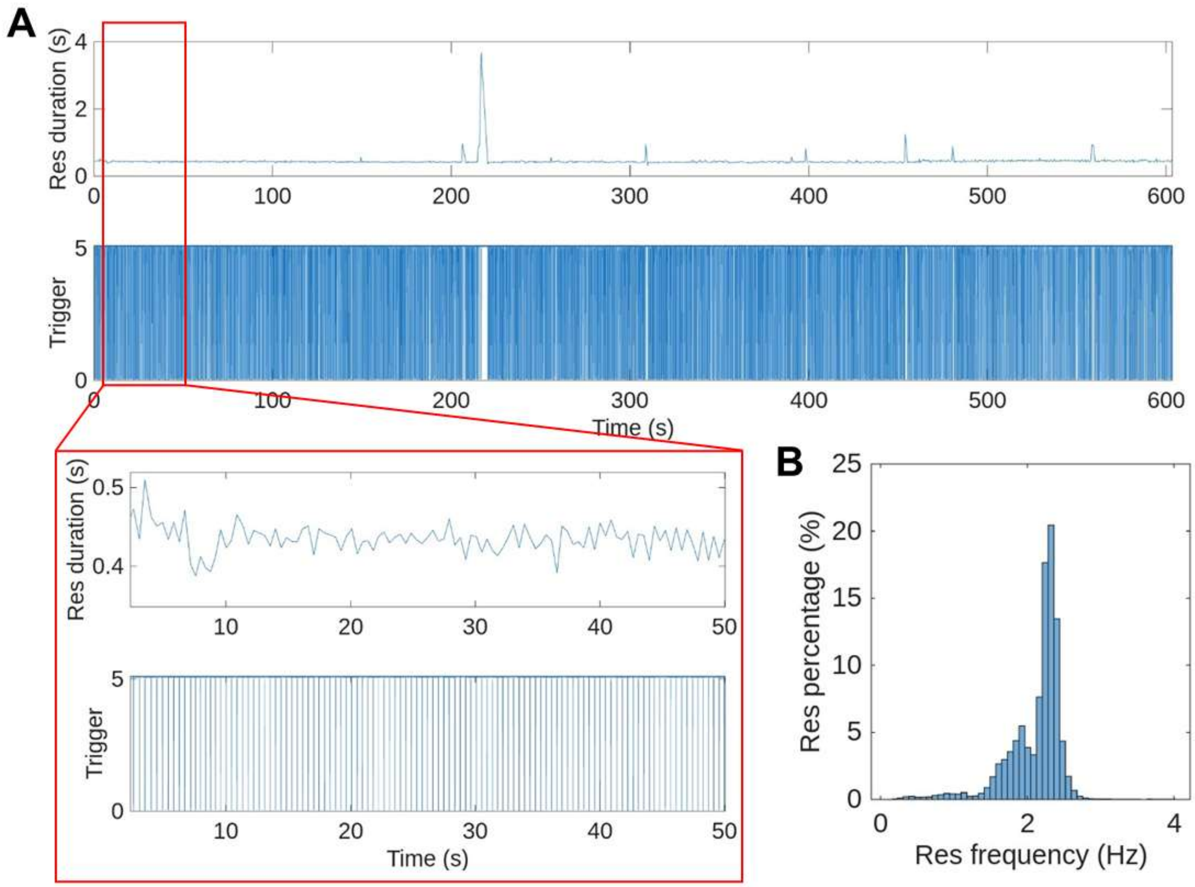


**Supplementary Figure 2. Respiration dynamics during NARS-fMRI acquisition in anesthetized mice.**

(A) Representative respiration trace recorded from an anesthetized mouse during NARS-fMRI acquisition. Enlarged images show the dynamic changes in respiration rate (i.e. the duration between two respiratory triggers) across the imaging session and relative to stimulation periods.

(B) Histogram of inter-breath intervals. The distribution highlights the shift in respiration rates during scanning, underscoring the need for respiration-gated NARS-fMRI acquisition with the ultrafast reshuffled k-t space trajectories.

**Supplementary Fig 3**

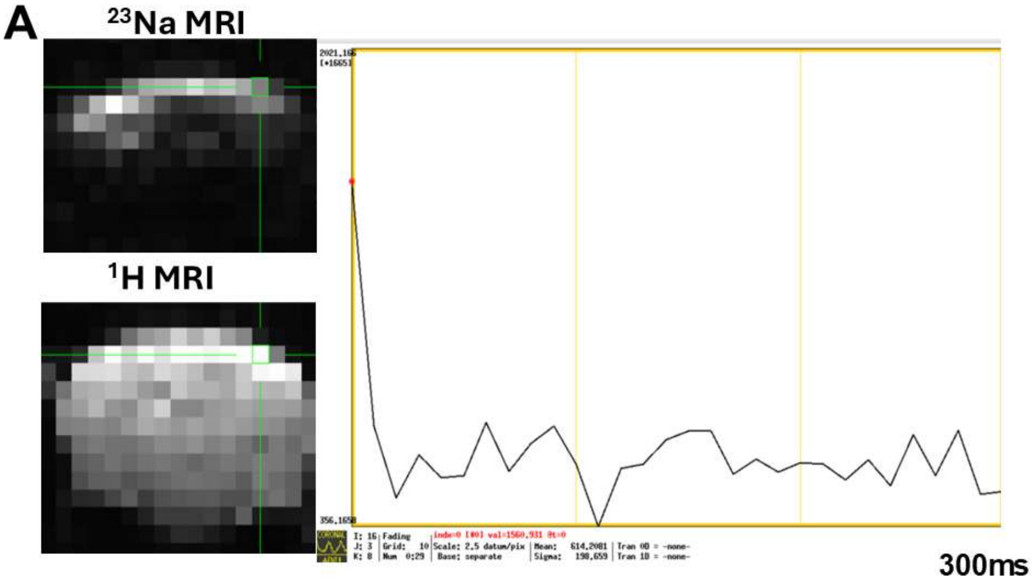


**Supplementary Figure 3. Recovery-dependent $^{23}$Na signal modulation across movie loops.**

**(A)** Mean $^{23}$Na signal intensity measured across 30 consecutive movie-loop time points spanning a 300 ms acquisition window during each stimulation epoch. The elevated signal observed at the first time point reflects reduced saturation of $^{23}$Na spins, arising from the longer respiration-dependent delay preceding the start of each stimulation epoch, which allows more complete longitudinal recovery before acquisition. Subsequent time points show progressively lower signal intensity as steady-state saturation is established across repeated ultrafast excitations.

### Supplementary Figure 4

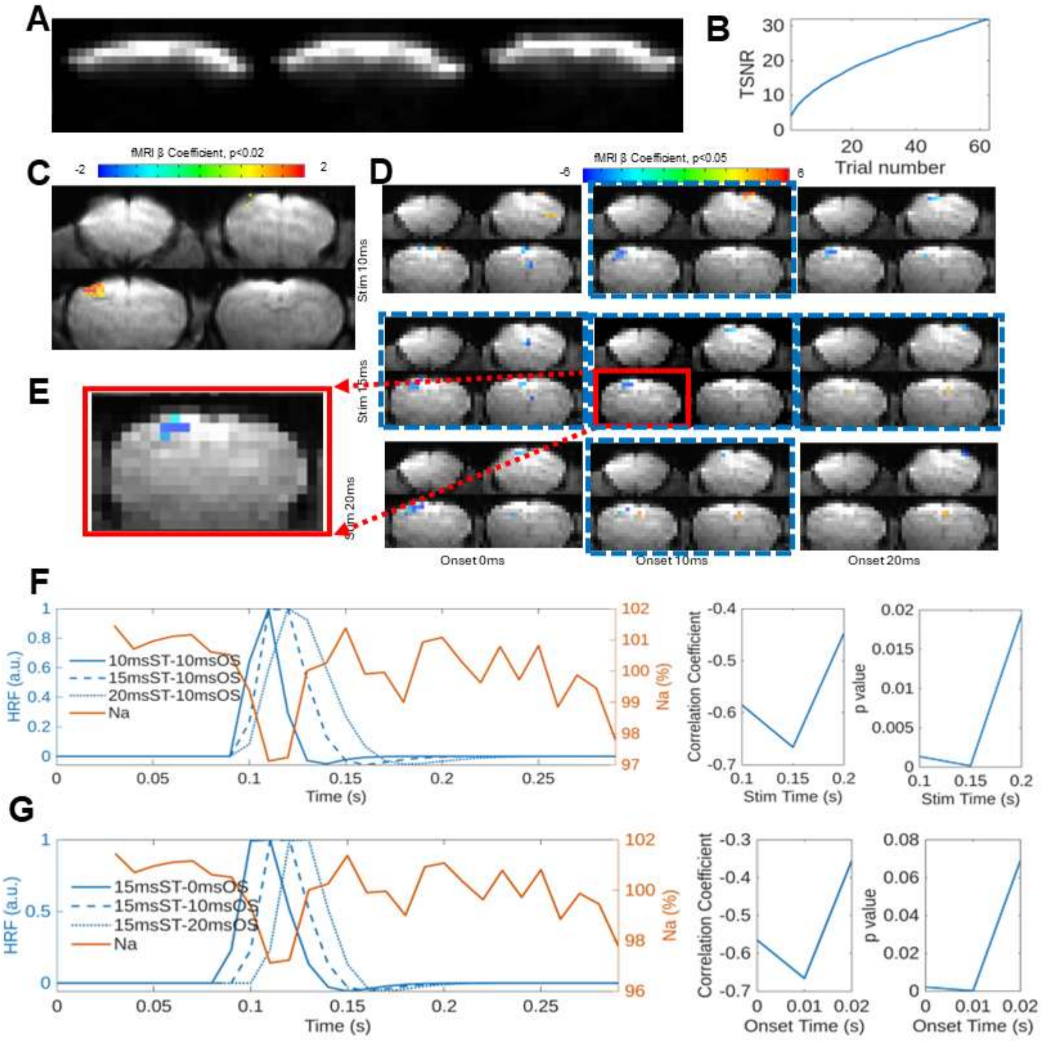


**Supplementary Figure 4. GAM-based ultrafast dynamic response function to model the millisecond NARS fMRI signal changes in rats.**

(A) Representative ultrashort-TE $^{23}$Na images (three adjacent slices) acquired with the reshuffled k-t space trajectory.

(B) Temporal SNR (TSNR) of the $^{23}$Na MRI time series as a function of trial number, showing progressive improvement with averaging across repetitions.

(C) BOLD fMRI statistical map (β coefficient; threshold $p < 0.02$) overlaid on the anatomical reference, showing focal activation in the FP-S1.

(D) Kernel-based mapping schemes: NARS statistical maps generated by correlating the ultrafast sodium series with brief gamma-like kernels spanning stimulus durations (rows: 10, 15, 20 ms) and onset shifts (columns: 0, 10, 20 ms). Maps are thresholded at $p < 0.05$ (color bar indicates β coeffiscient). Blue dashed boxes highlight parameter combinations yielding the most consistent FP-S1 maps.

(E) Zoomed view of the FP-S1 region (red box) to illustrate localization of the best-fit kernel map.

(F) Example sodium ROI time course (orange; $^{23}$Na signal, %) plotted together with candidate gamma-like kernels (blue) of different durations (10/15/20 ms) at a 10ms onset, illustrating the kernel family used for matching at native 10-ms sampling. Both correlation coefficients and corresponding p values versus kernel duration, suggesting the best fit for duration at 15ms.

(G) 23Na time course (orange) plotted with candidate kernels (blue) of different onset shifts at a 15ms duration (illustrating onset dependence). Correlation coefficients and corresponding p values versus kernel onset shift (onset time), showing the strongest fits for onsets within the 10 ms window.

### Supplementary Figure 5

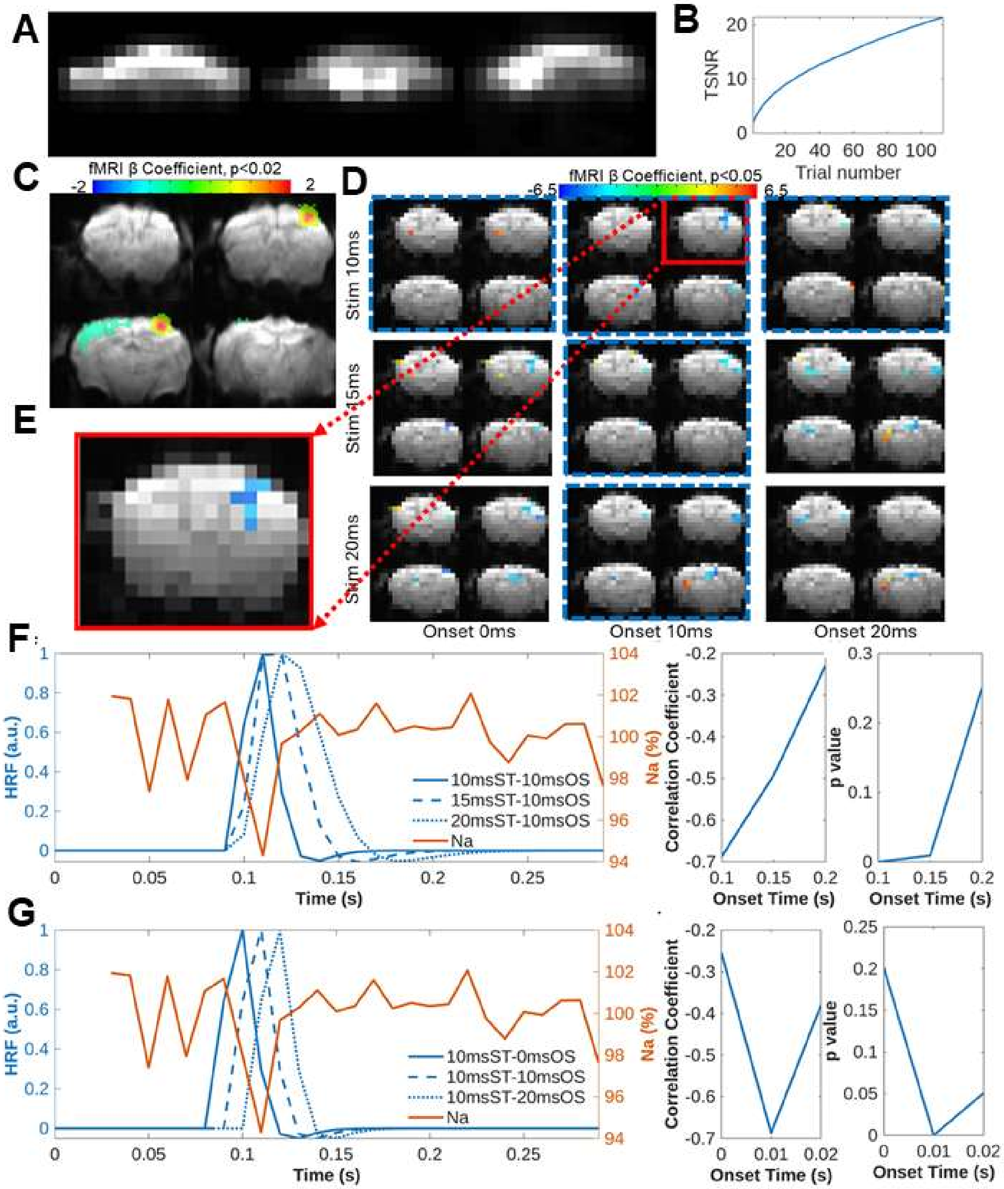


**Supplementary Figure 5. GAM-based ultrafast dynamic response function to model millisecond NARS-fMRI signal changes in mice.**

(A) Representative ultrashort-TE $^{23}Na$ images (three adjacent slices) acquired in mice using the reshuffled k–t space trajectory.

(B) Temporal SNR (TSNR) of the $^{23}Na$ MRI time series as a function of trial number, demonstrating progressive improvement with averaging across repetitions.

(C) BOLD fMRI statistical map (β coefficient; threshold $p < 0.02$) overlaid on the anatomical reference, showing focal activation in the forepaw somatosensory cortex (FP-S1).

(D) Kernel-based mapping schemes: NARS statistical maps generated by correlating the ultrafast sodium series with brief gamma-like kernels spanning stimulus durations (rows: 10, 15, 20 ms) and onset shifts (columns: 0, 10, 20 ms). Maps are thresholded at $p < 0.05$ (color bar indicates β coefficient). Blue dashed boxes highlight parameter combinations yielding the most consistent FP-S1 activation in mice.
(E) Zoomed view of the FP-S1 region (red box) illustrating localization of the best-fit kernel map.

(F) Example sodium ROI time course (orange; $^{23}$Na signal, %) plotted together with candidate gamma-like kernels (blue) of different durations (10/15/20 ms) at a 10 ms onset, illustrating kernel matching at the native 10 ms movie-loop sampling rate. Correlation coefficients and corresponding p values are plotted as a function of kernel duration, indicating a best fit at approximately 10 ms.

(G) Sodium ROI time course (orange) plotted with candidate kernels (blue) of different onset shifts at a fixed 10 ms duration, illustrating onset dependence. Correlation coefficients and corresponding p values plotted as a function of kernel onset shift show strongest fits for onsets within the first ~10 ms window.

**Supplementary Fig 6**

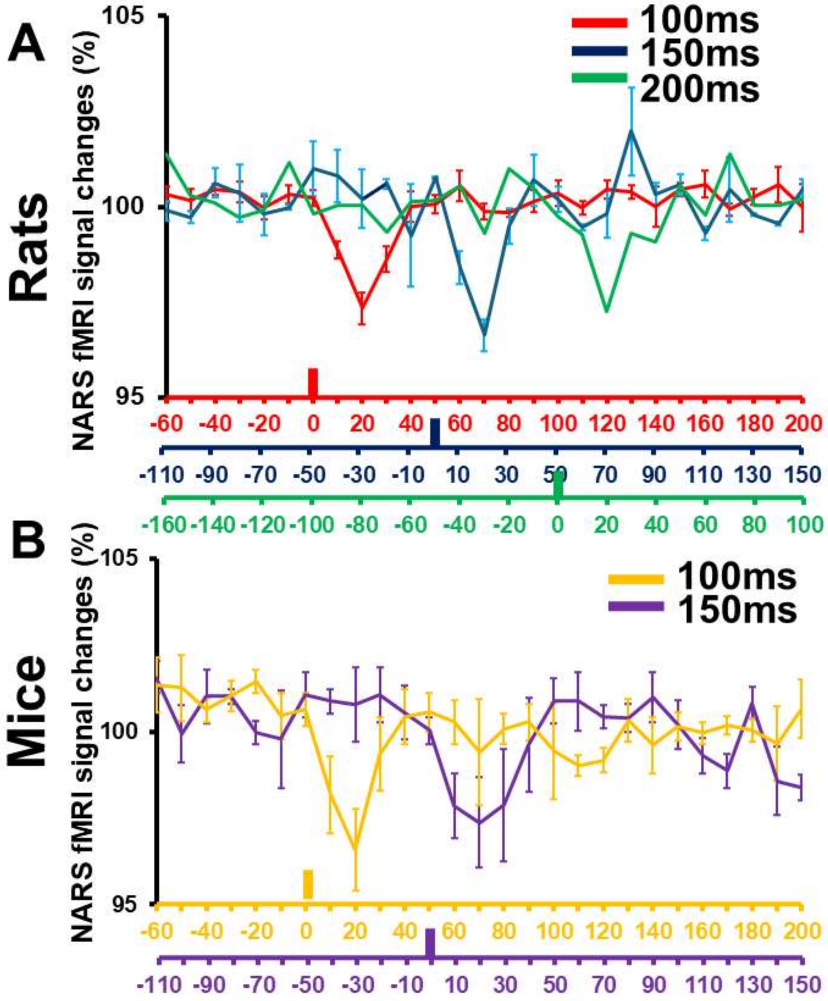


**Supplementary Figure 6. Stimulus onset-dependent millisecond timing of the NARS response in rats and mice.**

(A) Rat FP-S1 NARS-fMRI signal changes (%, mean ± s.e.m.) plotted at 10-ms movie-loop sampling rate for stimulation epochs in which the stimulus onset was varied within the 300-ms acquisition window. Traces are shown for 100 ms (red, n=5), 150 ms (blue, n=2), and 200 ms (green, n=1) stimulus onsets. Colored x-axes indicate the corresponding time bases after re-referencing each condition to its own stimulus timing.

(B) Same analysis for 100 ms (orange, n=5) and 150 ms (purple, n=4) stimulation of mice. The negative NARS deflection shifts with the imposed onset difference and remains locked to stimulus onset on the millisecond timescale, supporting timing specificity independent of the AFNI GLM.

**Supplementary Figure 7**

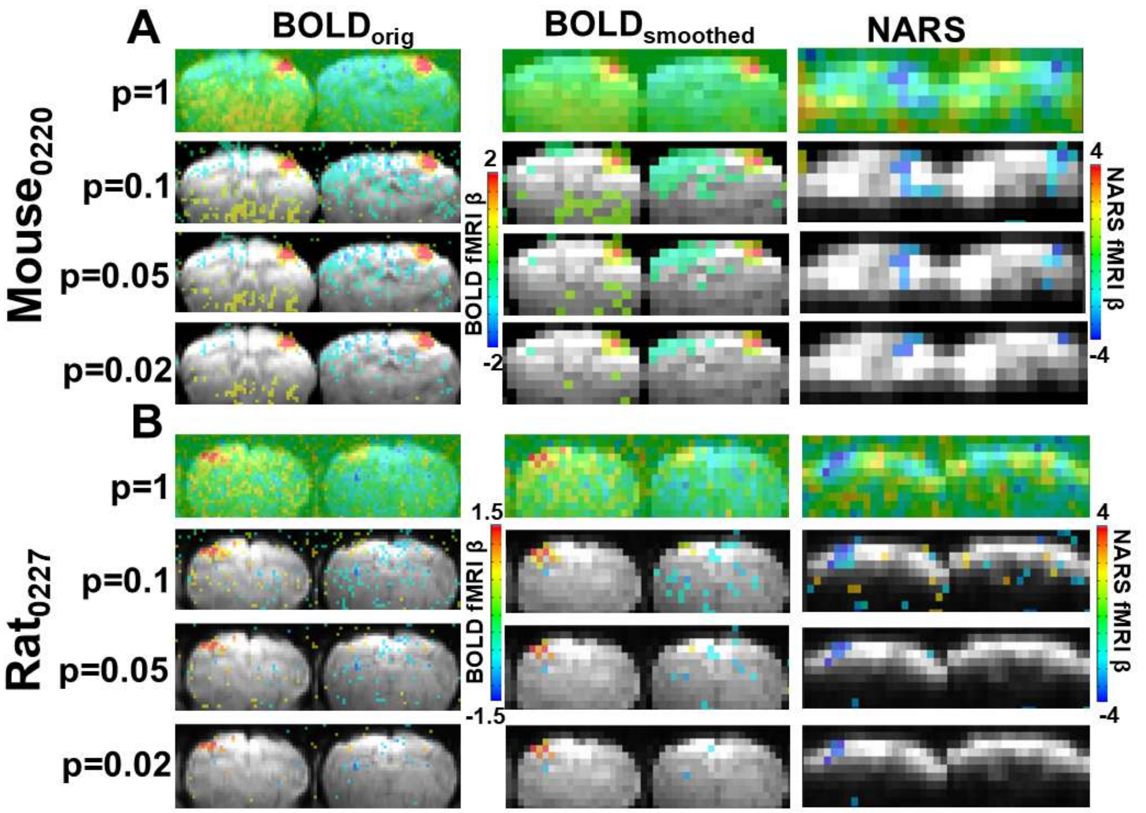


**Supplementary Fig. 7. Statistical threshold dependence of BOLD-fMRI and NARS-fMRI activation maps reveal robust focal FP-S1 responses beyond background noise.**

(A) Representative mouse datasets showing β-coefficient maps from BOLD-fMRI (original and spatially smoothed) and NARS-fMRI across progressively stricter statistical thresholds (p = 1, 0.1, 0.05, 0.02; derived from corresponding t-statistics). At permissive thresholds, widespread low-level fluctuations are observed, reflecting background noise contributions. Increasing statistical stringency progressively suppresses diffuse signals while preserving a focal activation pattern in the FP-S1. NARS-fMRI exhibits a localized negative $^{23}$Na response that spatially overlaps with the positive BOLD activation.

(B) Equivalent analysis in rat datasets demonstrates consistent behavior across species. Despite reduced background signals at stricter thresholds, focal FP-S1 activation persists in both BOLD- and NARS-fMRI maps. The stability of the localized NARS response under stringent thresholds supports a robust neuronal origin and argues against nonspecific coil sensitivity or sequence-related artifacts as primary drivers of the detected signal.

### Supplementary Figure 8

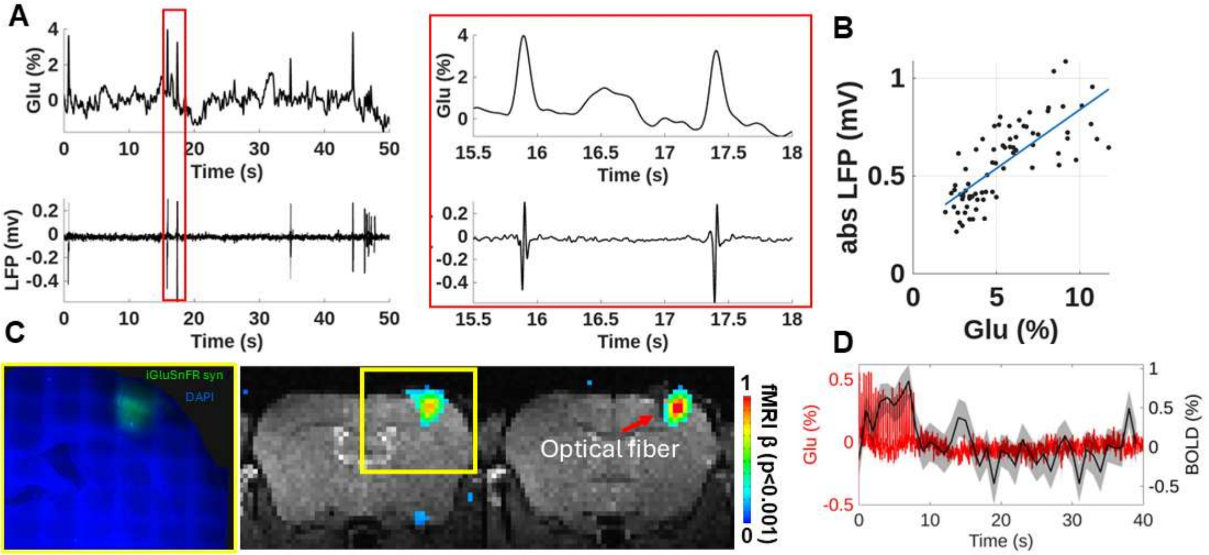


**Supplementary Figure 7. Characterization of the Glu with simultaneous LFP recording and BOLD fMRI.**

(A) Representative time courses of simultaneously recorded Glu (iGluSnFR, ΔF/F; green) and local field potential (LFP; black) signals from the FP-S1. Traces are aligned in time and show spontaneous transient events present in both modalities, illustrating their temporal correspondence at the single-event level.

(B) Scatter plot showing the relationship between peak amplitudes of spontaneous Glu transients and simultaneously recorded LFP events. Each point represents one spontaneous event detected in both modalities, with peak amplitudes extracted independently. Linear regression fitting curve demonstrates a positive amplitude-dependent coupling between Glu and LFP signals.

(C) BOLD fMRI statistical maps (threshold at $p < 0.001$) highlight the activation in FP-S1. The underlaid anatomical image indicates the optical-fiber location (arrow) relative to the activated region (insert, the representative fluorescence image showing iGluSnFR expression (green) with DAPI (blue) staining).

(D)The FP-S1 BOLD time courses (black) are plotted with the Glu signals (red). Glu recording exhibits rapid transients according to stimulus pulses (3Hz, 8s), whereas the BOLD response is delayed and broader, consistent with distinct neuronal versus hemodynamic dynamics.
